# Integration of phased Hi-C and molecular phenotype data to study genetic and epigenetic effects on chromatin looping

**DOI:** 10.1101/352682

**Authors:** William W. Greenwald, He Li, Paola Benaglio, David Jakubosky, Hiroko Matsui, Anthony Schmitt, Siddarth Selvaraj, Matteo D’Antonio, Agnieszka D’Antonio-Chronowska, Erin N. Smith, Kelly A. Frazer

## Abstract

While genetic variation at chromatin loops is relevant for human disease, the relationships between loop strength, genetics, gene expression, and epigenetics are unclear. Here, we quantitatively interrogate this relationship using Hi-C and molecular phenotype data across cell types and haplotypes. We find that chromatin loops consistently form across multiple cell types and quantitatively vary in strength, instead of exclusively forming within only one cell type. We show that large haplotype loop imbalance is primarily associated with imprinting and copy number variation, rather than genetically driven traits such as allele-specific expression. Finally, across cell types and haplotypes, we show that subtle changes in chromatin loop strength are associated with large differences in other molecular phenotypes, with a 2-fold change in looping corresponding to a 100-fold change in gene expression. Our study suggests that regulatory genetic variation could mediate its effects on gene expression through subtle modification of chromatin loop strength.

## Introduction

The three-dimensional (3D) architecture of the human genome is highly organized in the nucleus, bringing distant genomic regions into close spatial proximity and enabling colocalization of regulatory regions with their targets through chromatin looping^1–6^. Disease associated distal regulatory variation and expression quantitative trait loci (eQTLs) have been preferentially found at loop anchors^7–10^, and thus could potentially act by affecting chromatin looping. Previous studies examining the relationship between chromatin looping, cell types, genetic variation, and molecular phenotypes (i.e. gene expression, epigenetic variation) have suggested that cell type effects^2,11–13^ and genetic variation^5^ could cause large changes in chromatin looping, eliciting changes in molecular phenotypes. However, as chromatin structure has been shown to be an evolutionarily stable trait^14,15^, and genetic variation usually exerts subtle effects on molecular phenotypes^16^, regulatory genetic variants are *a priori* more likely to modulate loop strength than create or destroy loops. Quantitatively analyzing differential loop strength across cell types and between phased genetic variants (haplotypes) could elucidate whether genetic variation could modulate loop strength to the extent needed to affect molecular phenotypes. These analyses could provide insight into mechanisms underlying disease associated distal regulatory variation and help guide future studies aimed at understanding how genetic variants influence chromatin structure.

Previous studies that identified large changes in chromatin looping have been limited in that they did not differentiate between genetic effects and imprinting^2,5^, relied on targeted capture techniques, or examined only a handful of loci^5^. Chromatin loops within imprinted loci are commonly cited^2,5^ as examples of genetic effects on looping despite it being known that the allelic effects of imprinting are stronger and different from genetically driven allelic effects. The extent to which regulatory genetic variants - outside of imprinted loci - affect chromatin looping is therefore still unclear. Further, many targeted capture techniques (such as CTCF ChIA-PET^5^ and H3K27ac Hi-C ChIP^17^) simultaneously measure either regulatory region activity or protein binding with chromatin looping, and therefore could produce spurious functional associations by conflating regulatory activity differences with differences in chromatin loop strength.

Independent measurement of molecular phenotypes and chromatin looping via CHiP-seq and HiC, however, would enable the unbiased examination of the relationship between chromatin loop and epigenetic changes. A phased collection of Hi-C and molecular phenotype data across two cell types could therefore enable the study of the function of regulatory genetic and cell type effects on chromatin loops.

In this study, we generated a resource of phased, high resolution Hi-C chromatin maps from induced pluripotent stem cells (**iPSCs**) and iPSC-derived cardiomyocytes (**iPSC-CMs**) from seven individuals in a three-generation family, and phased previously published RNA-seq, H3K27ac ChIP-seq, and 50X WGS data from the same individuals. We identified chromatin loops, and quantitatively characterized cell type associated looping, finding that while loops tended to be present in both cell types, some loops exhibited significantly increased strength within one cell type. These cell type associated loops (**CTALs**) were more likely to harbor distal eQTLs, and were associated with the location and strength of differential molecular phenotypes. Additionally, we found that small magnitude changes in chromatin loops were proportionally associated with large changes in molecular phenotypes, with a 2-fold change in looping corresponding to a 100-fold change in gene expression. We found that imbalanced chromatin loops (**ICLs**) between haplotypes were not associated with large allelic imbalances in molecular phenotypes, and were primarily located in imprinted regions or associated with copy number variation; these results suggest that regulatory genetic variants are not associated with large changes in chromatin loop strength. Finally, despite observing smaller differences in loop strength across haplotypes than between cell types, we show that the relationship between loop strength and molecular phenotypes is consistent between cell types and haplotypes, suggesting that small loop differences are likely functionally relevant. Therefore, our study suggests that regulatory genetic variation could mediate its effects on gene expression through subtle modification of chromatin loop strength.

## Results

### Sample and data collection

Molecular data was obtained from iPSCs and their derived cardiomyocytes (iPSC-CMs) from seven individuals in a three-generation family from iPSCORE (the **iPSC** collection for Omics **RE**search)^18^ (Figure 1A, Table S1A). Fibroblasts from these seven individuals were reprogrammed using non-integrative Sendai virus vectors^19^, from which eleven iPSC lines were generated and subsequently differentiated into thirteen iPSC-CM samples using a monolayer-based protocol^20^. From the eleven iPSC and thirteen iPSC-CM samples, we generated chromatin interaction data via *in situ* Hi-C^2^. Additionally, from these and other iPSC and iPSC-CM samples from the same seven individuals, we integrated functional genomic data that was generated as part of a concurrent manuscript (RNA-seq, H3K27ac ChIP-seq, and ATAC-seq; Figure 1B and Table S1B; see methods) which also describes the differentiation efficiency and quality of all iPSC and iPSC-CM lines used in this study. Finally, we obtained single-nucleotide variants (**SNVs**) and somatic and inherited copy-number variants (**CNVs**) for the seven individuals from ~45X whole-genome sequencing (**WGS**) and genotype arrays from previously published work^18,21^.

**Figure 1.**
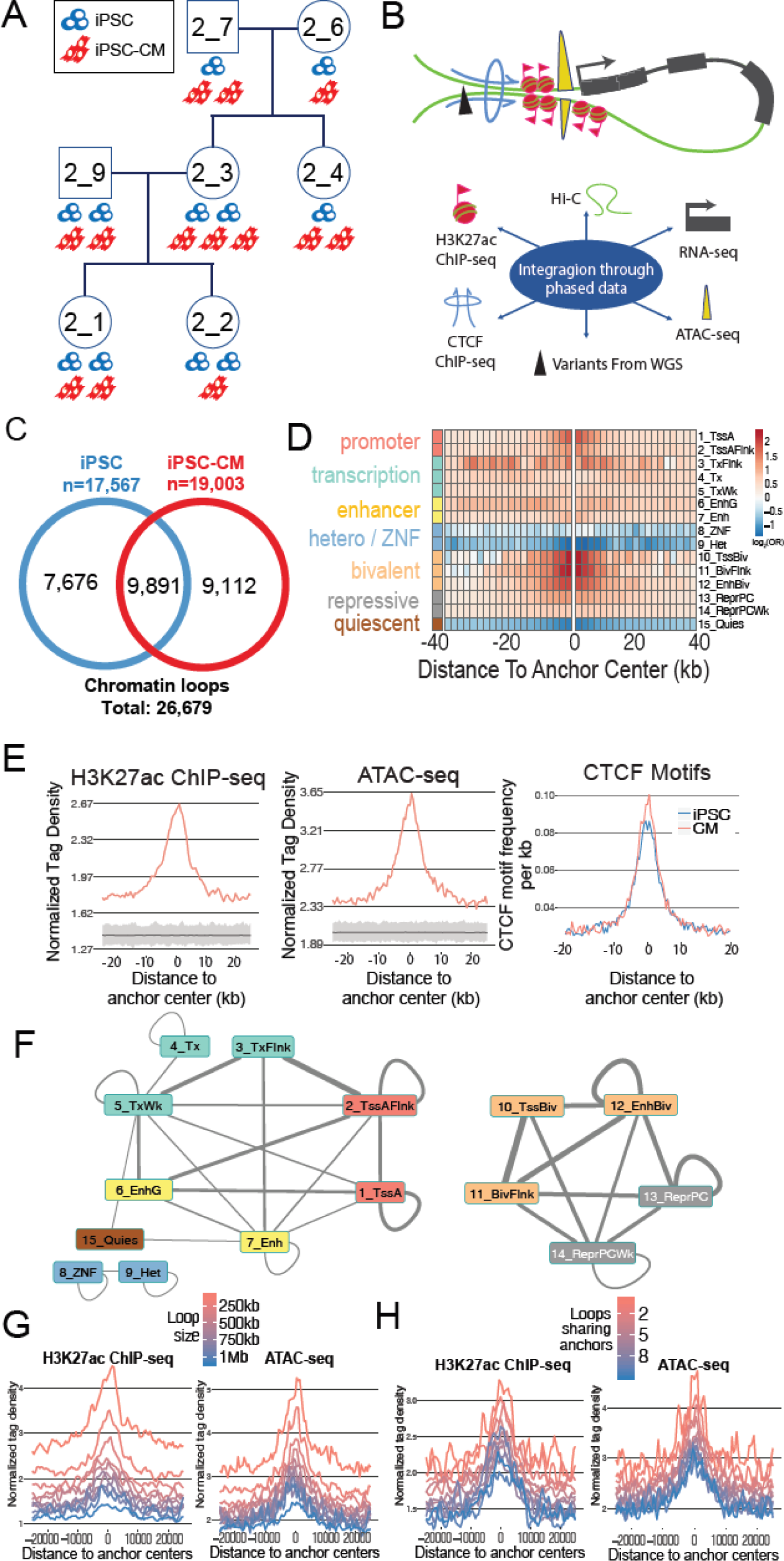
Chromatin contact maps and loops in iPSC and iPSC-CM. (A) Pedigree of the seven individuals used in this study. Cell icons below each subject indicate the number of iPSC lines and iPSC-CM samples used in the Hi-C experiments. iPSC lines are shown in blue, iPSC-CM samples are shown in red. (B) Schematic showing the data types used in this study depicting how they colocalize at loop anchors. (C) Venn diagrams showing the number of chromatin loops unique and common to both cell types. (D) Heatmap showing enrichment of regulatory regions near iPSC-CM called loop anchor centers. The 15 ROADMAP chromatin states of fetal heart tissue (E083) were used, and the log2 odds ratio of enrichment is indicated by color for each 2kb interval across an 80kb window. (E) Density plots showing distribution of epigenetic marks and motifs relative to the center of loop anchors. Normalized tag densities from H3K27ac ChIP-seq (left) and ATAC-seq (middle) are shown for loops called in iPSC-CM. Grey regions below the peak signals indicate the results from 1,000 null loop sets. CTCF motif frequency per kb (right) is shown for loops called in iPSCs (blue) or iPSC-CMs (red). (F) Network diagram showing two discrete subnetworks of fetal-heart chromatin states at iPSC-CM called loops, with edges connecting statistically significant pairs of chromatin states found at opposing. The thickness of the edge indicates the odds ratio of significance, and the presence or absence of an edge indicates statistical significance. (G-H) Density plots showing distribution of epigenetic marks and motifs relative to the center of loop anchors stratified by loop size (G) or loop complexity (H). Average normalized tag densities from H3K27ac ChIP-seq (left) or ATAC-seq (right) around loop anchors are shown.

### Identification and characterization of chromatin loops in iPSCs and iPSC-CMs

We characterized the 3D chromatin structure of iPSCs and iPSC-CMs by identifying chromatin loops in each cell type genome wide. From the *in situ* Hi-C data, we obtained 1.74 billion long-range (≥20kb) intra-chromosomal contacts after aligning and filtering ~6 billion Hi-C read pairs across all twenty-four Hi-C samples (Figure S1A, Table S1C). We performed hierarchical clustering of the contact frequencies by cell type across individuals and observed high correlations within each cell type (Figure S1B). We therefore pooled the data within each cell type to create the highest resolution chromatin maps in iPSCs and iPSC-CMs (or any other iPSC derived cell type) to date (~2kb map resolution; Figure S1C). As loop calling algorithms often identify distinct loops, and are dependent on the resolution parameters specified for their analysis^22^, we called chromatin loops from these maps utilizing two algorithms (HICCUPS and Fit-Hi-C) at multiple resolutions, identifying 17,567 loops in iPSCs (**iPSC called loops**), and 19,003 iPSC-CM loops (**iPSC-CM called loops;** Tables S1D-S1E). We examined the overlap of the loops between cell types and found that 37.1% of the total 26,679 loops were called in both cell types (Figure 1C), which is consistent with previous work showing differential detection of loops between cell types, but is lower than the overlap found when comparing high resolution maps to those of lower resolution^2^. These data comprise the largest loop set from Hi-C data to date, and provide a resource for the analysis of long range gene regulation across the genome.

To establish that the iPSC and iPSC-CM called loops showed functional properties consistent with their identified cell type, we examined the distribution of H3K27ac and ATAC peaks, CTCF motifs, and chromatin states (iPSC for iPSCs; fetal heart for iPSC-CMs, the most epigenetically similar cell type) near loop anchors. In both cell types, we found enrichments for active and bivalent chromatin states (Figure. 1D & S1D), H3K27ac (Figure 1E left & S1E), chromatin accessibility (Figure 1E middle & S1F), and CTCF motifs at loop anchors (Figure 1E right). We next examined the types of chromatin states that were statistically significantly paired together (Fisher’s Exact p < 0.05) and found two subnetworks, one with active chromatin states and the other with repressed or bivalent chromatin, which were discrete in iPSC-CMs (Figure 1F) and crossed over through the bivalent states in iPSCs (Figure S1G). These results indicate that the identified chromatin loops include those with active regulatory interactions (e.g. promoter-enhancer interactions), those with repressive interactions (e.g. polycomb complexes), and those with other types of chromatin states at their anchors. As differentiation can be accompanied by an increase in interactions among repressed regions^12^, we next examined two physical characteristics that may reflect higher order structure: the distance between the anchors of a loop (**loop size**), and the number of distinct loops sharing an anchor with a loop (**loop complexity**).

We tested whether these physical characteristics were associated with functional characteristics and observed that both loop size and loop complexity were associated with H3K27ac and ATAC-seq signals (Figure. 1G & 1H), with larger loops and more complex loops tending to be repressed with lower functional activity, and smaller loops and less complex loops tending to be active with high functional activity. Overall these analyses established the iPSC called loop set and iPSC-CM called loop set for further analysis.

### Quantification of differential chromatin looping between cell types

To determine if the chromatin loops called in only one of the cell types were specific to that cell type, or whether they were also present in the other cell type but not called, we performed a quantitative comparison of loop intensity between the cell types. For all loops, identified in either one or both cell types, we compared the total normalized read count intensity (log2 counts per million, **logCPM**) of the interactions between both cell types. We observed that the majority of loops that were called in both cell types (grey in Figure 2A) had high logCPMs in both cell types, whereas the loops that were only called in a single cell type (blue or red in Figure 2A) tended to have overall low logCPMs and often showed highly similar contact intensities between cell types. We did not observe, however, loops with a high logCPM in one cell type, and a very low logCPM in the other. These results indicate that chromatin loops that were differentially called between cell types were often of low logCPM intensity, and were therefore likely to be inconsistently identified by the loop calling algorithms, and that the differences in loops between cell types were not due to the establishment of novel loops in only one cell type. We therefore identified loops that showed quantitative differences between iPSCs and iPSC-CMs by comparing normalized read counts across cell types at each loop identified in either cell type (edgeR glmQLFit q < 0.01; Figure 2B). This analysis resulted in four loop sets (Table S1F): 1) all loops called in any cell type (**union loop set**, total: 26,679), 2) loops with statistically higher intensity in iPSCs (iPSC cell type associated loops; **iPSC-CTALs**, total 2,906), 3) loops with statistically higher intensity in iPSC-CMs (**CM-CTALs**, total 2,915), and 4) loops that were not statistically significantly different between the two cell types (**non-CTALs**, total 20,858).

**Figure 2:**
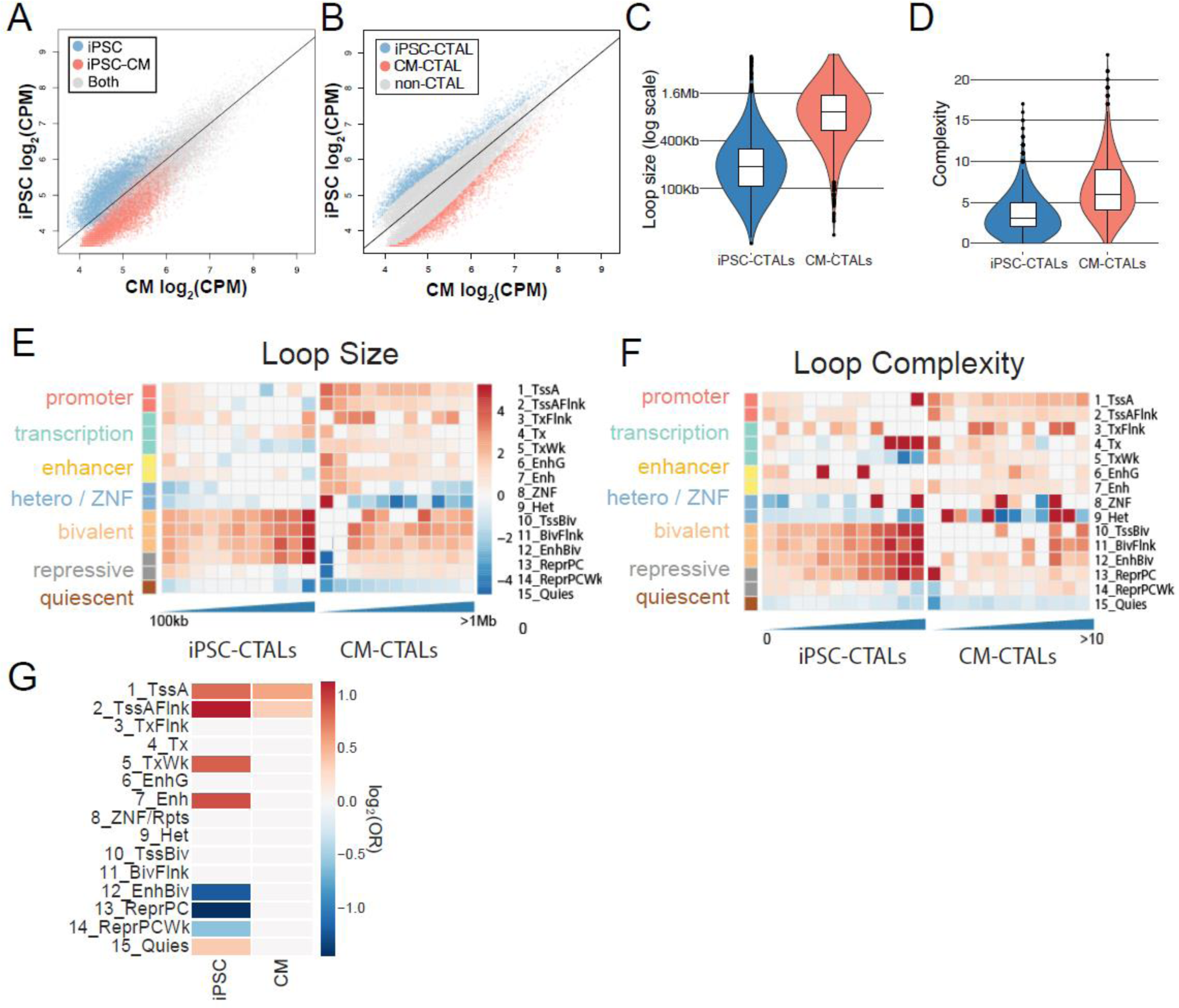
Differential chromatin states and sizes in CTALs recapitulate changes in looping across differentiation. (A-B) Scatterplots showing contact frequency in counts per million (CPM) of all loops identified in either iPSCs or iPSC-CMs. The solid black lines indicate the function y = x. (A) Points are colored to indicate loops called in only iPSC (blue), or iPSC-CM (red), or both (gray). (B) Points are colored to indicate loops with significantly increased loop strength in iPSCs (iPSC-CTAL; blue), iPSC-CMs (CM-CTAL; red), or neither (non-CTAL; gray). (C-D) Violin plots showing distributions of loop size (C), and loop complexity (D) for CTALs. (E-F) Heatmap showing enrichment of regulatory regions near iPSC-CTAL (left) and CM-CTAL (right) at loop anchor centers with loops stratified by (E) size or (F) complexity. The 15 ROADMAP chromatin states of iPSC (E020) or fetal heart tissue (E083) were used, and the log2 odds ratio of enrichment is indicated by color. CTALs broken down by size into 100kb windows (E), or complexity (F). (G) Heatmap of log2(odds ratio) from a Fisher’s exact tests for enrichments of differential chromatin states across CTAL anchors. White cells indicate a nonsignificant Fisher’s Exact test (FDR q > 0.05)

### CTALs are associated with regulatory changes during differentiation

Throughout differentiation, chromatin architecture has been reported to specialize and become more cell type specific^12,13,23^. To determine whether quantitative differences in loop strength were associated with functional and physical changes associated with differentiation, we examined the physical and regulatory characteristics of CTALs. We observed that CM-CTALs were overall significantly larger (Mann-Whitney p < 2.2×10^−16^; Figure 2C) and more complex (Mann-Whitney p < 2.2×10^−16^; Figure 2D) than iPSC-CTALs. Additionally, we found active chromatin states to be preferentially enriched at smaller (Figure 2E) and less complex (Figure 2F) loops. Next, we examined whether CTALs for each cell type were enriched for overlapping cell type-specific regulatory regions (Figure 2G). For example, to test whether iPSC-CTALs were more likely to harbor an iPSC-specific active promoter, we restricted the analysis to loops overlapping an iPSC active promoter and tested whether the proportion of loops overlapping an iPSC specific active promoter was higher within iPSC-CTALs than non-iPSC-CTALs. We found iPSC-CTAL and CM-CTAL anchors to be enriched for differential active promoters, and iPSC-CTAL anchors to be enriched for differential active enhancers (Figure 2G). These enrichments suggest that CTALs capture cell type specific chromatin dynamics. We also observed that iPSC-CTAL anchors which overlapped iPSC bivalent chromatin to be more likely to overlap fetal heart bivalent chromatin. Overall, these findings show that CTALs were enriched for cell type specific functional and regulatory regions, and indicate that iPSC-CM differentiation is associated with larger and more complex loops, consistent with increased polycomb repressed looping^12^.

### Functional characterization of CTALs

To analyze the functionality of CTALs, we examined the relationship between loop strength and eQTLs, differential gene expression, and differential epigenetics across cell types. We first examined whether loops which colocalize iPSC-eQTLs (previously identified from a cohort including these individuals^21^) to their eGenes had stronger contact intensities within iPSCs that iPSC-CMs. We found a strong enrichment (Mann Whitney-U p ~ 1×10^−293^) for iPSC contact intensity above non eQTL-eGene loops (Figure 3A), indicating that loops with higher strength in a cell type may be more likely to harbor functional genetic variation. Next, we examined whether differential molecular phenotypes were preferentially located at CTAL anchors. We identified differential H3K27ac peaks and genes using ChIP-seq and RNA-seq data generated from iPSC and iPSC-CM samples from the same seven individuals (see methods). We obtained a total of 23,570 differential H3K27ac peaks (DE peaks) and 5,307 differential genes (DE genes) between iPSCs and iPSC-CMs. We found that DE genes and DE peaks were preferentially located at CTAL anchors (Fisher’s exact p < 0.05, Figure 3B) compared to the union loop set. Together, these results suggest that cell type associated loop strength is relevant for other cell type associated molecular phenotypes.

**Figure 3.**
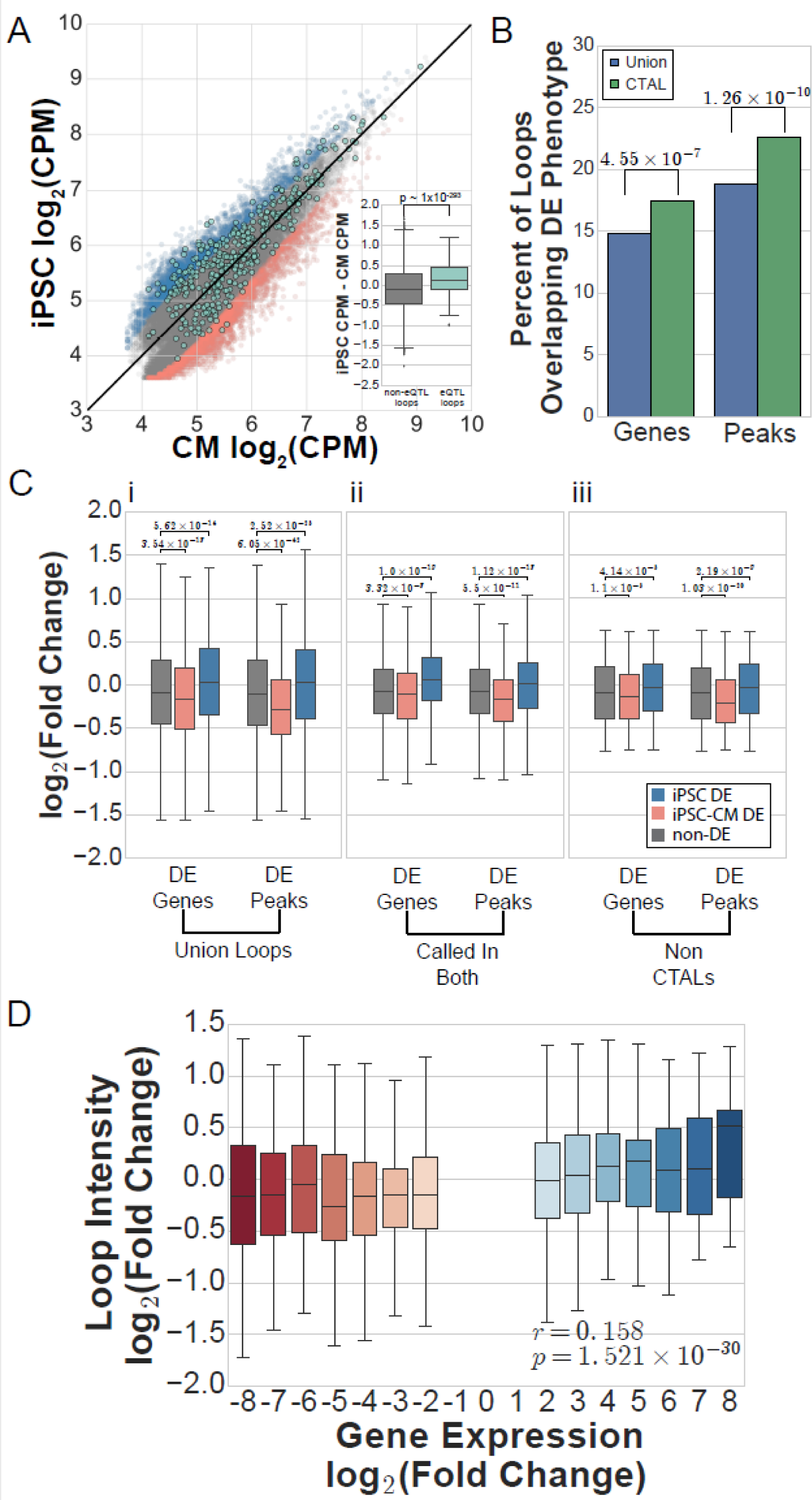
Quantitative variation in chromatin loops is associated with differential gene expression and H3K27ac across cell types. (A) Scatterplot showing iPSC vs. iPSC-CM contact frequencies in counts per million (CPM) for all union loops. The black line indicates the y = x function. Background points indicate iPSC-CTALs (blue), non-CTALs (grey), and CM-CTALs (red). Overlaid on this are points indicating iPSC eQTL-eGene containing loops (teal). The boxplot in the lower right corner of the scatter plot shows the difference between iPSC and iPSC-CM CPMs at each non-eQTL loop (grey) or eQTL loop (teal). Positive values indicate a loop has higher CPM in iPSCs, and negative values indicate a loop has higher CPM in iPSC-CM. The p-value was calculated from a Mann-Whitney U test. (B) Barplot showing the percent of CTALs (green) or union loops (blue) which overlap differentially expressed genes or H3K27ac peaks. P-values were found via a Fisher’s Exact test for the underlying counts of differentially expressed genes or peaks between union loops and CTALs. (C) Boxplots of the log2(fold change) of contact intensity at chromatin loops, with positive indicating strong in iPSCs and negative indicating stronger in iPSC-CMs, for all loops (i), loops called in both cell types (ii), or non-CTALs (iii) with anchors overlapping differentially expressed genes or H3K27ac peaks with higher expression or counts in iPSCs (blue), higher expression or counts in iPSC-CMs (red), or not overlapping a DE gene or peak (grey). P-values were found via a Mann-Whitney U test. (D) Boxplot showing the log2(fold change) of chromatin loop intensity for chromatin loops overlapping a differentially expressed gene, binned by the log2(fold change) of the gene. For both expression and chromatin looping, positive indicates stronger counts in iPSCs, and negative indicates stronger counts in iPSC-CMs. The Pearson correlation and p-value shown were calculated on the raw underlying data.

Next, we examined the quantitative association between loop strength and differential expression or H3K27ac across cell types. We tested whether the fold change in contact intensity across cell types was in the direction of the cell type with higher differential expression or H3K27ac. We found that across the union loop set, anchors overlapping DE genes expressed higher in iPSCs had significantly higher contact intensities in iPSCs, while anchors overlapping DE genes expressed higher in iPSC-CMs had significantly higher contact intensities in iPSC-CMs; similar patterns were found for DE H3K27ac peaks (Mann-Whitney-U p < 0.05; Figure 3C i). To establish that this association was due to differences in loop strength, rather than being driven by loops that were differentially called between the two cell types, we examined whether this association was still present within loops that were called in both cell types (i.e. the intersection of iPSC-CM and iPSC called loops). We found that the statistically increased contact intensity (Mann-Whitney-U p < 0.05) in the upregulated cell type remained within this set of loops, though the extent of the differences in chromatin looping were smaller (Figure 3C ii). Thus, we next examined whether these differences could be observed at non-CTALs (i.e. loops with nonsignificant differences across cell types) and found that these loops were still significantly stronger in the expected direction when they overlapped a DE molecular phenotype at their anchor (Figure 3C iii). These results suggest that subtle variation in chromatin looping across cell types may be functional. Finally, to examine whether chromatin loops proportionally varied with the strength of gene expression differences between cell types, we examined the correlation between fold changes in gene expression and chromatin looping at loops with anchors overlapping promoters of differentially expressed genes (Figure 3D). We observed a significant correlation (r = 0.158, p < 1.6×10^−30^) between the two phenotypes; however, the magnitudes at which the phenotypes varied were quite different, with gene expression varying up to 250-fold, and chromatin looping varying less than 3-fold. Overall, these results indicate that small magnitude changes in chromatin looping may be functional as they are associated with large magnitude changes in gene expression.

### Haplotype-based interrogation of chromatin loops, gene regulation, and gene expression

To enable the functional characterization of haplotype-specific chromatin looping, we phased the Hi-C, H3K27ac, and RNA-seq data to obtain haplotype-associated phenotype data (Tables S2-S4). We first phased the WGS genotype data for these seven individuals using a combination of Hi-C-based phasing and family structure, resulting in an average of 2.01M phased heterozygous variants per individual (Figure S2A-C, see methods). Next, we assigned informative reads from H3K27ac and RNA expression to each individual’s maternal or paternal haplotype using an established method^24^, and then identified significant ASE peaks or genes (FDR q < 0.05) within each individual using a binomial test. We identified a total of 189 ASE peaks (mean 43 per individual) in iPSCs and 618 ASE peaks (mean 119 per individual) in iPSC-CMs, and 2,582 ASE genes (mean 647 per individual) in iPSCs and 2,214 ASE genes (mean 503 per individual) in iPSC-CMs.

To characterize haplotype-specific chromatin looping, we identified significantly imbalanced chromatin loops (**ICLs**) genome wide. Within each cell type, we assigned informative Hi-C contacts carrying a phased allele to each haplotype (Figure 4A) and examined allelic imbalance across all loops. For each individual, we identified imbalance via a binomial test on a half normal distribution (see methods), following which we combined the p-values across individuals with Fisher’s method. This process identified 54 total ICLs: 27 from iPSCs, and 27 from iPSC-CMs. Across the 54 ICLs, we observed similar maternal allele ratios in both cell types within each individual which were highly correlated (0.73 < Pearson’s r < 0.97; example individuals in Figure 4B&C; all individuals in Figure S3A-B) which suggests that, while loop imbalance was consistent across both cell types, we were limited in statistical detection of the imbalance due to the sparsity of HiC data. Therefore, to increase power for these analyses, for each of the 26,679 chromatin loops in the union set, we pooled contacts for each individual across their corresponding iPSCs and iPSC-CMs. We observed a median of 50 informative contacts per individual per loop, which corresponds to 100% power to identify ICLs with an allelic imbalance ratio of 70% or higher with α = 0.02 in an individual (Figure S3C), or at α = 2×10^−5^ when all samples display similar imbalance and are combined with Fisher’s method meta-analysis. Within each subject, a mean of 6.08% of all chromatin loops showed significant imbalance at p < 0.05 (binomial test on a half normal distribution; see methods), slightly higher than the statistically expected 5% by chance; however, only a mean of 0.1% (26.6) were significant under FDR q < 0.05 in each individual (Figure 4D). To identify ICLs which were consistently imbalanced across individuals, we again combined associations using a Fisher’s method meta-analysis for each loop, and identified 7.49% of chromatin loops as ICLs at p < 0.05, indicating that consistent allelic imbalance occurs more frequently than by chance; however, only 114 ICLs were significant after multiple testing corrections at FDR q < 0.05 (equivalent to p < 2×10^−5^) even with the combined cell type data, and the majority of these loops had small allelic differences (Figure 4E). These results indicate that chromatin loops mainly exhibit subtle differences across haplotypes, as across cell types, and suggest that large haplotype differences chromatin looping occur infrequently.

**Figure 4.**
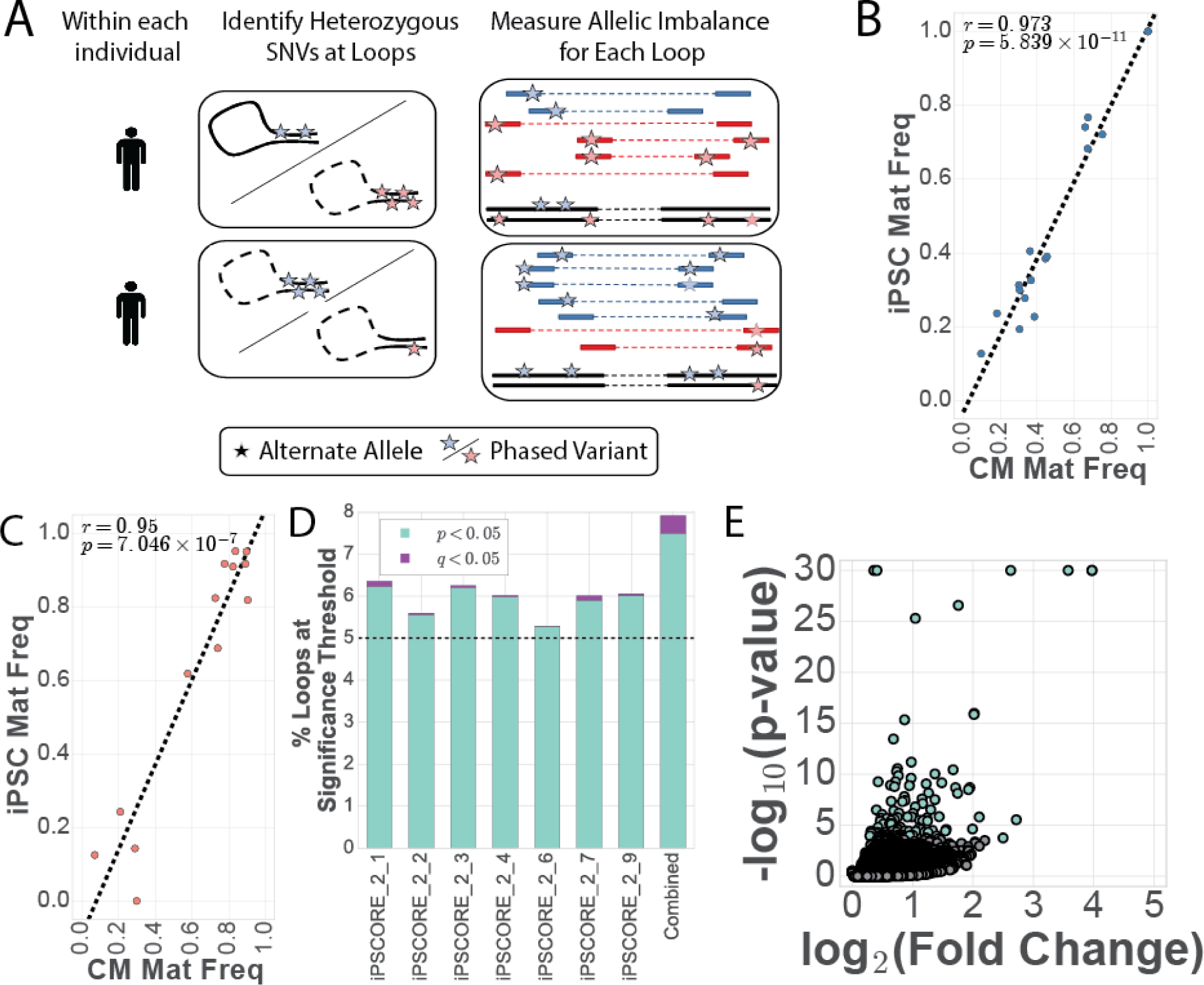
Identification of haplotypic differences of chromatin conformation. (A) Schematic showing approach to quantify chromatin loop imbalance within each individual. Examples for two different individuals are shown. Variants were phased using Hi-C and family structure (see methods), and each contact was assigned to its corresponding haplotype based on the phase of heterozygous SNVs it contained. (B-C) Scatter plot showing comparison between iPSC and iPSC-CM maternal haplotype frequencies for one of the seven individuals at ICLs identified in either (B) iPSCs or (C) iPSC-CMs. Linear regression correlation and p-value are reported for each cell type. Similar plots for all 7 subjects are in Figure S3. (D) Barplot showing the percent of loops associated with haplotypic imbalance at p < 0.05 shown in teal, with those also q < 0.05 shown in purple. Bars are shown for each individual separately (left 7 bars), or for the results of a Fisher’s method meta-analysis p-value (combined; right most bar). A dashed line is drawn at 5% to indicate the number of ICLs expected by chance to be significant at p < 0.05. (E) Volcano plot showing the log10(p-value) vs the log2(fold change) for each loop with the combined data. Significant points (ICLs) are shown in teal.

### ICLs are associated with imprinting and CNVs

We next examined whether the 114 genome-wide significant haplotype-specific chromatin loops (i.e. ICLs) were statistically more likely to also be cell type specific loops (i.e. CTALs), or overlap genomic features previously shown to be associated with differential chromatin looping (imprinted genes^2,5^ and somatic and inherited CNVs^25,26^). We hypothesized that chromatin loops that were variable across cell types may be more variable in general, and thus ICLs would be more likely to be CTALs. We compared the proportion of ICLs that were also iPSC-CTALs, CM-CTALs, iPSC called, or iPSC-CM called loops to the corresponding proportion of union loops. However, we found no significant differences for any association (p > 0.05 for all tests; Figure 5A), indicating that loops which varied between haplotypes were not more likely to vary between cell types. We next compared the distribution of genomic features known to cause large allelic differences within ICLs and the union loop set (Figure 5B). We observed that, compared to the union loop set, ICLs were statistically more likely to contain imprinted genes (ICL: 10.5%; all: 2.7%; Fisher’s exact p = 5.8×10^−5^), and somatic (ICL: 7.0%, all: 1.0%; Fisher’s exact p = 1.8×10^−5^) and inherited (ICL: 27.2%, all: 18.3%; Fisher’s exact p = 2.03×10^−2^) CNVs previously identified in these samples^21^. To examine whether these trends held across all levels of imbalance significance, we quantified the extent of association of each genomic feature with chromatin loop allelic imbalance as a function of ICL p-value. For imprinted genes, as the p-value threshold increased, the odds ratio increased almost log-linearly, whereas CNV overlap increased, but to a lesser extent (Figure 5C). These results suggest that genetic imprinting, and to a lesser extent CNVs, may be a strong driver of allelic imbalanced chromatin looping.

**Figure 5.**
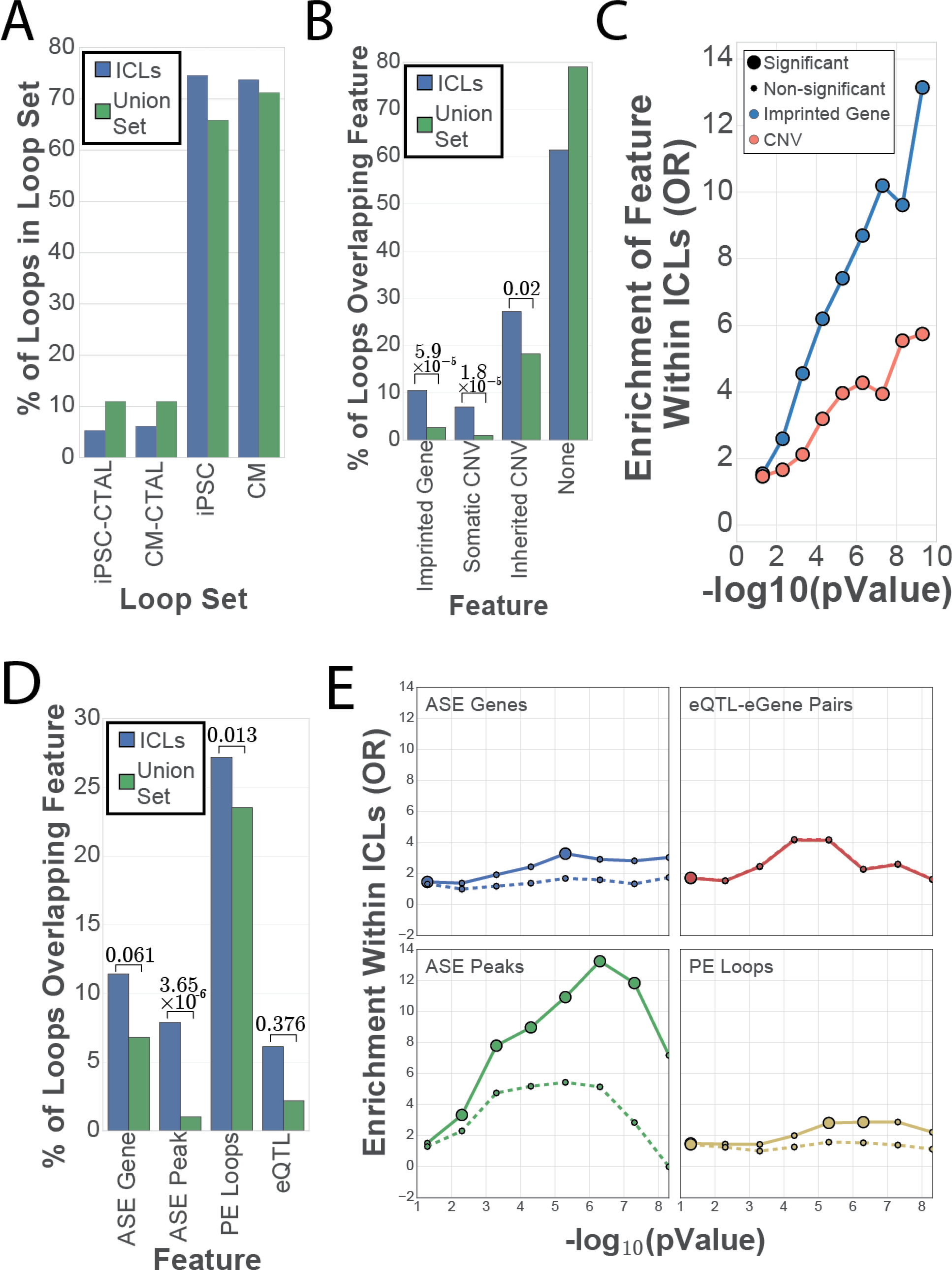
Functional characterization of haplotypic differences in chromatin conformation. (A) Barplot showing the percent of union loops (green) or ICLs (blue) contained within each loop-set. (B) Barplot showing the percent of union loops (green) or ICLs (blue) containing the given genomic feature within it (i.e. the genomic feature overlapped the region between the start of the first anchor and the end of the second anchor). P-values were found via a Fisher’s exact test. (C) Line plot showing odds ratio from a Fisher’s exact test for ICL enrichment above the union set for containing an imprinted gene (blue) or containing either an inherited or somatic CNV (red) as a function of the -log_10_ of the ICL imbalance p-value. Large circles indicate that the test was significant after Bonferroni correction, and small circles indicate a non-significant association. (D) Barplot showing the percent of union loops (green) or ICLs (blue) overlapping the given genomic feature at an anchor. P-values were found via a Fisher’s exact test. (E) Line plot showing odds ratio from a Fisher’s exact test for ICL enrichment above the union set for containing the labelled feature as a function of the -log_10_ of the ICL imbalance p-value, for either all loops (solid lines), or loops that do not contain an imprinted gene or CNV (dashed lines). Large circles indicate that the test was significant after Bonferroni correction, and small circles indicate a non-significant association.

We next examined whether ICLs were enriched for functional allele-specific differences at their anchors by quantifying the enrichment for containing an ASE gene or ASE H3K27ac peak at their anchors, or being a promoter-enhancer or eQTL-eGene loop. We found ASE peaks to be enriched at ICL anchors and being a promoter enhancer loop to be enriched (Fisher’s Exact p < 0.05; Figure 5D); notably, despite the increased percentage of eQTL-eGene loops in ICLs, as only 7 eQTL-eGene loops were ICLs (585 eQTL-eGene loops in total), this increase was nonsignificant. To determine whether regulatory genetic variation was associated with these differences, we excluded the effects from imprinting and CNVs, and examined these associations across a range of imbalance thresholds (Figure 5E). The removal of imprinted regions and CNVs greatly attenuated the association and resulted in a loss of significance for the two molecular phenotypes, and PE loop status, over almost all ranges of imbalance significance. These results suggest chromatin loops vary across haplotypes much more subtly (i.e. allelic ratio <70%) than gene expression or H3K27ac, and where variation is larger, it is mainly driven by imprinting and/or CNVs rather than genetic variation. Overall, these results show that large allelic imbalances in molecular phenotypes are restricted to chromatin loops primarily located in imprinted regions or associated with copy number variation, and that regulatory genetic variants are not associated with large changes in chromatin loop strength.

### Small quantitative differences in looping are associated with large differences in molecular phenotypes

We observed that differential loop strength was associated with differential molecular phenotype strength across cell types (Figure 3), but not across haplotypes (Figure 5). To resolve this discrepancy, we compared the relationship of loop strength and molecular phenotype strength across haplotypes to the same relationship across cell types. We first compared the general variability of chromatin loops outside of imprinted regions and CNVs across cell types (Figure 6A) to the differences across haplotypes (Figure 6B). We found that more chromatin loops varied to a larger degree across cell types than across haplotypes, with ~35% of loops exhibiting a log2 fold change of 0.5 (1.4-fold) or higher across cell types, but only ~5% across haplotypes (Figure 6C), consistent with haplotype associated differences being considerably smaller than cell type associated differences. We therefore next examined the association between chromatin loop strength and gene expression. Across cell types and haplotypes, we found a positive and significant correlation between gene expression fold change and chromatin loop fold change; notably, we found large changes in gene expression to be associated with small changes in chromatin looping (Figure 6D). The slope of the association between haplotypes was also similar to that observed across cell types, with a 2-fold change in chromatin loop strength corresponding to a 100-fold change in gene expression. We next examined this same relationship between loop strength and H3K27ac strength, and found significant associations across cell types and haplotypes, though the fold changes between H2K27ac and looping were more similar (β=0.06) than that of gene expression and looping (β =0.02). This consistency and magnitude difference indicate that large differences in gene expression, and moderate changes in H3K27ac, are associated with small differences in chromatin looping, and suggest that small changes in chromatin looping are likely functionally relevant. Additionally, as the association between gene expression and loop strength was consistent across haplotypes, these results suggest that genetic variation could exert effects on gene expression through small modifications of loop strength. Overall, these results suggest that genetic variation could mediate its effects on gene expression by subtly modifying chromatin loop strength, as small changes in looping were associated with large changes in molecular phenotypes.

**Figure 6:**
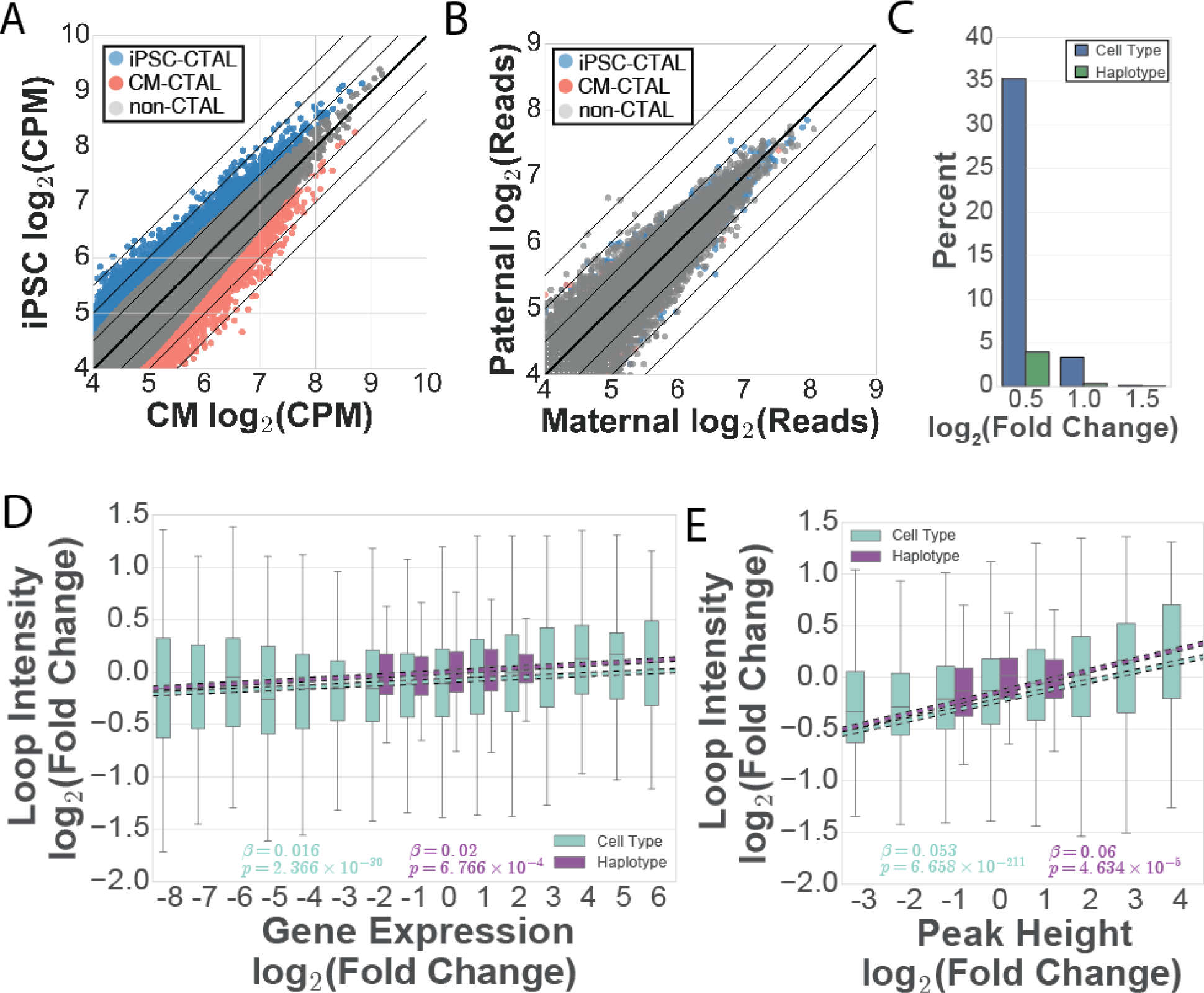
Comparison of chromatin loop, gene expression, and H3K27ac variability across cell types and haplotypes. (A-B) Scatterplots showing (A) contact frequency in counts per million (CPM) across cell types or (B) read counts across haplotypes of all union loops colored by CTAL status. The solid bold lines indicate the function y = x, and other lines indicate absolute fold changes of log2(0.5), log2(1), and log2(1.5). (C) Percent of loops with at least the shown log2(Fold Change) or across cell types (blue) or haplotypes (green). (D-E) Boxplot showing the log2(fold change) of chromatin loop intensity for chromatin loops overlapping a (D) differentially expressed or ASE gene, or (E) differential or ASE H3K27ac peak, binned by the log2(fold change) of the (D) gene or (E) peak. Boxes are shown for cell type comparisons in teal, and haplotype comparisons in purple, linear regressions are plotted with dashed lines, and beta’s and p-values are shown and colored from the raw data in each data set independently. For all data, positive fold change indicates stronger counts in iPSCs, and negative fold change indicates stronger counts in iPSC-CMs.

## Discussion

Here, we provide a resource of phased molecular phenotype data in two cell types for seven individuals who are a part of a three-generation family, and use this data to perform an in depth, genome-wide, functional examination of changes in chromatin loop strength across cell types and haplotypes. We find that chromatin loops, including those associated with differential molecular phenotypes, are not uniquely present in a single cell type or on a single haplotype, but instead show quantitative differences. We then show that, across cell types and haplotypes, changes in chromatin loops are associated with proportionally similar changes in gene expression and the epigenome, though the magnitude of changes are quite different, with a 2-fold change in looping corresponding to a 100-fold change in expression and a 33-fold change in H3K27ac. The depth of Hi-C reads at a given chromatin loop measures spatial proximity (i.e. the distance in 3D space between two anchors), but is biased by factors dependent on the loop’s genomic coordinates (number of restriction enzyme sites near the anchors, anchor GC content, and mapping uniqueness)^27^. These factors are held constant between cell types or haplotypes for a given loop; our observed differences in loop strength are therefore likely to reflect true differences in spatial proximity. Our work therefore suggests that small cell-type or haplotype associated quantitative differences in spatial proximity may be associated with large changes in gene expression and the epigenome. Additionally, while the phased data we provide is a resource for future studies on the function of regulatory variation, the Hi-C maps are the highest resolution maps for iPSCs and iPSC-CMs currently available and thus will be an important resource for the prioritization of functional variants and their potential gene targets in these cell types.

Unlike previous studies^2,5^, we stratified our haplotype analyses by whether the genomic regions were known to undergo imprinting. We found that allelic imbalance at chromatin loops over 70% was strongly enriched for imprinted regions. While the allelic imbalance outside of imprinted regions were small in magnitude, they were correlated with large changes in other molecular phenotypes, suggesting that the regulatory genetic variants at these loci could exert subtle changes in loop strength that may be responsible for alteration of gene expression. The identification of these causal variants, however, is likely to be challenging as we found chromatin loops to show very minor deviations from allelic imbalance of 50%, even in the case of large differences in associated molecular phenotypes. For example, for a gene with an ASE of 98%, the expected chromatin loop imbalance would be ~52%, which would require ~2,000X coverage to obtain an uncorrected p-value of 0.05. Therefore, it may be practically infeasible to identify specific variants that alter chromatin structure through the genome-wide identification of chromatin loop QTLs with Hi-C. For the validation of specific variants, future studies seeking to study whether genetic variants are associated with chromatin loop imbalance should consider using an unbiased targeted loop capture assay with higher sensitivity than Hi-C.

Finally, our work provides some insight into the ongoing question of whether changes in chromatin looping cause changes in gene expression, or if changes in gene expression cause changes in looping^1,2,11–13,15,17,28,29^. It has been established that the creation of new chromatin loops can alter gene expression^30^, however is has been less clear whether altering gene expression results in meaningful changes in chromatin loops^11,12,31^. Evaluating whether chromatin loop changes are meaningful requires an understanding of the scale at which functional changes in chromatin loops occur. As our findings suggest that subtle changes are functional, we believe these discordant interpretations could have arisen from studies either not being sufficiently powered to detect small effects, or from discounting small changes as nonfunctional. Our work therefore provides a foundation for future studies to quantitatively examine how changes in looping elicit changes in expression (or vice versa) and suggests that studies designed to detect small magnitude changes in chromatin loop variability may be needed to delineate the relationship between chromatin loop imbalance and gene expression.

## Author Contributions

Conceptualization, W.W.G., H.L., E.N.S., and K.A.F.; Methodology, W.W.G., E.N.S., and K.A.F.; Software, W.W.G., H.L., and H.M.; Validation, H.L.; Formal Analysis, W.W.G., H.L., P.B., M.D., and D.J.; Investigation, W.W.G., H.L., A.D.A., P.B., A.S., and S.S.; Data Curation, W.W.G., H.M.; Writing - Original Draft, W.W.G., H.L., E.N.S., and K.A.F.; Writing – Review & Editing, W.W.G., E.N.S., and K.A.F.; Visualization, W.W.G., and H.L; Supervision, E.N.S., and K.A.F.; Project Administration, W.W.G, E.N.S., and K.A.F.; Funding Acquisition W.W.G., P B., D.J., A S., S.S., E.N.S., and K.A.F.

## Acknowledgments

This work was supported in part by a California Institute for Regenerative Medicine (CIRM) grant GC1R-06673 and NIH grants HG008118-01, HL107442-05, DK105541-03 and DK112155-01. RNA-seq were performed at the UCSD IGM Genomics Center with support from NIH grant P30CA023100. WWG was supported by the National Heart, Lung, And Blood Institute of the National Institutes of Health under Award Number F31HL142151. DJ was supported by the National Library of Medicine Training Grants T15LM011271. PB is supported in part by the Swiss National Science Foundation Postdoc Mobility fellowships P2LAP3-155105 and P300PA-167612. Whole genome sequencing was performed at Human Longevity, Inc. Arima Genomics was supported by NIH grantR41HG008118. We would like to acknowledge Yunjiang Qiu for initial scripts and help with HiC loop calling.

## Data Availability

All genomic data will be available through dbGAP accessions phs000924 (Hi-C, RNA-seq, CHiP-seq, ATAC-seq) and phs001325 (whole genome sequence SNV and CNV genotypes).

## Methods

### Subject enrollment

The seven individuals used in this study were recruited as part of the iPSCORE project^18^. This recruitment was approved by the Institutional Review Boards of the University of California, San Diego and The Salk Institute (Project no. 110776ZF), and consent forms were received from each subject. Subject information including sex, age, and ethnicity were collected during recruitment (Table S1A). Skin biopsy was performed to obtain fibroblasts for iPSC reprogramming, and blood samples were collected for whole genome sequencing.

### iPSC derivation, iPSC-CM differentiation, and sample collection

Cell line derivation and differentiation were performed as described in Benaglio *et al.* [in review]. From the seven individuals, fibroblast samples from skin biopsies were reprogrammed using non-integrative Cytotune Sendai virus (Life Technologies)^19^ following the manufacturer’s protocol. Each independent reprogramming resulted in one or more iPSC clones of the subject.

At passages 12-13, genomic integrity of at least one iPSC clone per subject was assessed using Illumina HumanCoreExome arrays, and pluripotency of iPSCs was assessed for most clones in this study by flow cytometry of the pluripotency markers SSEA4 and TRA-1-81^18^. iPSCs of each clone were harvested between passages 12 to 40, resulting in a total of 38 iPSC samples used in this study (Table S1B). Each iPSC clone was then used to generate multiple independent iPSC-CM differentiations using a monolayer protocol^20^, resulting in a total of 27 iPSC-CM samples used in this study. Among these iPSC-CM samples, 11 of them were subjected to purification via 4 mM Sodium L-Lactate at Day 15 after the start of differentiation and collected at Day 25^32^; one iPSC-CM sample was subjected to lactate purification at Day 11 and collected at Day 16; the rest of the iPSC-CM samples were not subjected to lactate purification and collected at Day 15 (Table S1B). Across all molecular assays detailed below, lactate purified and non-lactate purified iPSC-CM samples showed similar profiles; we therefore combined data across the two protocols. Single-nucleotide variants (**SNVs**) and copy-number variants (**CNVs**) of these individuals were obtained from ~40X whole-genome sequencing (**WGS**) results described in iPSCORE (dbGaP id: phs001325.v1.p1) and by DeBoever et. al.^21^.

### Hi-C data generation

For each of the 11 iPSC and 13 iPSC-CM Hi-C samples, we performed *in situ* Hi-C on 2-5 million cells. Hi-C libraries were prepared using *in situ* Hi-C as previously described^2^. Briefly, cells were crosslinked at a final concentration of 1% formaldehyde and quenched using 200 mM glycine. Crosslinked cells were then lysed and nuclei were digested with 100U Mbol overnight at 37°C. Next, fragmented ends were biotinylated for 90min at 37°C, and the sample was diluted and proximity ligated for 4 hours at room temperature. Crosslinks were reversed by the addition of SDS, ProteinaseK, and NaCl, and allowed to incubate overnight at 68°C. Samples were then purified by ethanol precipitation, resuspended in 100uL 1X Elution Buffer, fragmented using a Covaris S2 instrument, and size selected using AmpureXP beads. Subsequently, biotinylated ligation junctions were pulled down using T1 Streptavidin beads. Hi-C libraries were prepared using streptavidin beads by performing end-repair, dA-tailing, and adapter ligation, following which PCR amplification and purification was performed. The resulting libraries were sequenced on an Illumina HiSeq 4000 machine to obtain 150bp paired-end reads.

### RNA-Seq data generation

RNA-seq data was obtained from the Benaglio *et. al* [in review] manuscript. Specifically, total RNA was isolated using the Qiagen RNAeasy Mini Kit from frozen RTL plus pellets, including on-column DNAse treatment step. RNA was eluted in 60 μ! RNAse-free water and run on a Bioanalyzer (Agilent) to determine integrity. Concentration was measured by Nanodrop. Illumina Truseq Stranded mRNA libraries were prepared and sequenced on HiSeq2500, to an average of 40 M 100 bp paired-end reads per sample. RNA-Seq reads were aligned using STAR^33^ with a splice junction database built from the Gencode v19 gene annotation^34^. Transcript and gene-based expression values were quantified using the RSEM package (1.2.20)^35^ and normalized to transcript per million bp (TPM).

### ChIP-Seq data generation and peak calling

We used the H3K27ac data published in Benaglio *et. al* [in review]. For H3K27ac, 2 x 10^6^ fixed cells were lysed in 60 μl of MAGnify™ Chromatin Immunoprecipitation System Lysis Buffer (Thermo Scientific) and sonicated using Bioruptor 200 (Diagenode) for 35-45 min of 30 sec on/30 sec off cycles. H3K27ac antibodies (Abcam ab4729, lots GR183922-2 (1.75 μg) or GR184333-2 (1 μg)) were coupled for 2 hours to ProteinG Dynabeads (Thermo Scientific), and used for overnight chromatin immunoprecipitation in IP buffer (1% Triton-X, 0.1% DOC, 1x TE, 1x Roche Complete Proteinase Inhibitor tablets (RCPI)). Beads were washed five times with washing buffer (50 mM Hepes pH 8, 1% NP-40, 0.7% DOC, 0.5M LiCl, 1mM EDTA and 1x RCPI) and once with TE buffer. For transcription factors, 1-2 x 10^7^ cells were lysed in 300 μl RIPA buffer (1xPBS, 1% NP-40, 0.5% DOC, 0.1% SDS, RCPI) and sonicated for 70-80 min with instrument and setting as above. Five μg of SRF antibody (Santa Cruz Biotechnology, sc-335x, lot I1014) were incubated with Dynabeads for 2 hours and washed with BSA 0.5% in PBS. Chromatin was diluted to 1 ml of RIPA buffer and added to the beads for overnight IP. Five washes were performed with washing buffer (100 mM Tris pH 7.5, 500 mM LiCl, 1% NP-40, 1% DOC and 1x RCPI), followed by one wash with TE. DNA was eluted and reverse crosslinked overnight in elution buffer (10 mM Tris-HCl pH 8, 1 mM EDTA, 1% SDS) at 65°C. DNA was purified using Qiagen MinElute PCR Purification kit, quantified by Qubit (Thermo Scientific) and submitted to library preparation and barcoding using KAPA Hyper Library preparation kit (KAPA Biosystems). Libraries were sequenced on an Illumina HiSeq2500 or a HiSeq4000 to an average of 35 M 100 bp paired-end reads per sample.

ChIP-Seq reads were mapped to the hg19 reference using BWA54. Duplicate reads, reads mapping to blacklisted regions from ENCODE, reads not mapping to chromosomes chr1-chr22, chrX, chrY, and read-pairs with mapping quality Q <30 were filtered. Peak calling was performed using MACS2^36^ (‘macs2 callpeak -f BAMPE -g hs -B –SPMR –verbose 3 –cutoff-analysis –call-summits -q 0.01’) using pooled BAM from all iPSC or iPSC-CM samples for each ChIP-Seq antibody and with reads derived from sonicated chromatin not subjected to IP (i.e. input chromatin) from a pool of samples used as a negative control.

### ATAC-Seq data generation and peak calling

We used the data published in Benaglio *et. al* [in review]. Specifically, the ATAC-Seq protocol has been adapted from Buenrostro et al.^37^. Frozen nuclear pellets of 5 x 10^4^ cells each were thawed on ice, suspended in 50 μL transposition reaction mix (2.5 μL Tn5 transposase in 1x TD buffer, Illumina Cat# FC-121-1030), and incubated for 30 min at 37°C. Reactions were purified using Qiagen MinElute kit, eluted in 10 μL water and amplified using the KAPA real-time library amplification kit (KAPA Biosystems) with barcoded adaptors. PCR reactions were terminated after 10 to 13 cycles and purified using AmPure XP beads (Beckman Coulter). Samples were size selected using SPRIselect beads (Beckman Coulter) to a size range of 150 to 850 kbp and sequenced on an Illumina HiSeq2500 to an average depth of 30 M 100 bp paired end reads.

ATAC-Seq reads were aligned using STAR to hg19 and filtered using the same protocol as for ChIP-Seq. In addition, to restrict the analysis to regions spanning only one nucleosome, we required an insert size no larger than 140 bp, as we observed that this improved sensitivity to call peaks and reduced noise. Peak calling was performed using MACS2 on merged BAM files of iPSC and iPSC-CM meta-samples with the command ‘macs2 callpeak –nomodel –nolambda – keep-dup all –call-summits -f BAMPE -g hs’, and peaks were filtered by enrichment score (q < 0.01).

### Creation and analysis of Hi-C contact maps

For each sample, Hi-C reads were first aligned to human reference genome hg19 using BWA-MEM (version 0.7.15)^38^ with default parameters. Forward and reverse reads from the paired-end data were aligned independently to allow for identification of split reads that represent ligations between two genomic loci due to spatial proximity^2^. Paired-end reads were then reconstructed, processed, and filtered using the Juicer pipeline^39^, resulting in the removal of: unmapped reads, abnormal split reads (split reads that cause ambiguous positioning of the contact), read pairs within the same restriction enzyme fragment, low mapping quality read pairs (MAPQ < 30), and duplicate reads. Subsequently, read pairs that were less than 2kb apart were removed to avoid self-ligated fragments. These filtered read pairs (contacts) were subsequently used to generate chromatin contact maps for each sample via Juicer. To create Hi-C contact maps on a per individual basis, contacts were pooled across all samples of a particular cell type for each individual, and to create maps of iPSC and iPSC-CM, contacts were pooled across individuals within the respective cell type. These processes resulted in a set of binary .hic files, which were utilized to obtain raw and Knight-Ruiz (KR)^40^ normalized counts as well as normalization vectors of contact frequency matrices via Juicebox command line tools^41^ at various resolutions used throughout this study.

### Correlation of Hi-C contact maps between samples

The KR normalized contact matrices of each sample were retrieved from the .hic files at various resolutions (100kb, 250kb, 500kb, and 1Mb) using Juicebox^41^. The contact matrices were then vectorized in order to calculate Pearson’s correlation between each of the samples in R. Hierarchical clustering analyses of the Pearson’s correlation were performed in R using hclust with default settings and (1-Pearson’s correlation) as dissimilarity height.

### Identification of chromatin loops

Chromatin loops in iPSC and iPSC-CM were called using both Fit-Hi-C^42^ and HICCUPS^2,41^ as summarized in Figure S4A. For Fit-Hi-C, loops were called in meta-fragment resolutions that each contained a fixed number of consecutive restriction enzyme (RE) fragments, ranging from 10 to 30 RE fragments. Loop calling procedures for each resolution are summarized in Figure S4B. First, significant interactions (FDR q < 0.01) were identified through jointly modeling the contact probability using raw contact frequencies and KR normalization vectors with the Fit-Hi-C algorithm (Step 1). Next output of Fit-Hi-C was pruned by requiring that: 1) the interaction itself was significant; 2) for each anchor of the interaction, 3 of the 5 immediately upstream or downstream bins from the opposing anchor were significant (Step 2). We then merged high-confidence interactions within 20kb using pgltools^43^ (Step 3), discarded interactions that did not have any other interactions within 20kb, and retained the most significant call at each interaction event (Step 4).

For HICCUPS, loops were called using fixed-size bin resolutions from 5kb to 25kb at 1kb bin size intervals using parameters summarized in Table S1G. Briefly, default parameters of peak size (p) and window size (i) were used to call loops at 5kb and 10kb resolutions provided by HICCUPS^41^. For 6kb, 7kb, 8kb, and 9kb resolutions, the values of these two parameters were interpolated from the 5kb and 10kb values, and rounded to the closest integer. For resolutions greater than 10kb, the default 10kb parameters were used. Following loop calling, as performed by Rao & Huntley et al.^2^, for resolutions from 5kb to 10kb, loops within 20kb were merged using pgltools. For resolutions above 10kb, loops within twice the size of the anchor were merged using pgltools. At each merging event, the loop call with the most statistical significance provided from HICCUPS output was retained.

We next intersected loop calls across all resolutions within each calling method, retaining the highest-resolution call at each intersection event, and filtered loops to loops present in 3 HICCUPS resolutions, or 7 Fit-Hi-C resolutions, as these loops visually appeared to best represent the underlying Hi-C data (Figure S4C). We found a large number of loops that overlapped between Fit-Hi-C and HICCUPS (Figure S4D); however, many loops were unique to only one caller (Figure S4E). We therefore intersected the loops across calling methods, retaining the loop with the smallest total anchor size at each intersection event (Figure S4F), resulting in iPSC called and iPSC-CM called loop sets.

### Creation of the union loop set

To create the union loop set, we used pgltools merge to find all loops from the iPSC call set and iPSC-CM call set with both anchors within 20kb. This process led to merge events of 1, 2, or 3 loops, which were resolved as follows: 1) if there was only 1 loop present within 20kb (ie, only 1 loop set had a call), this loop was retained, 2) if there were 2 loops present within 20kb, the loops were merged by pgltools merge, 3) if there were 3 loops present, pgltools closest was used to identify which two loops were closest together; these two loops were merged, and the third loop was retained as its original call.

### Identification of cell type associated loops (CTALs)

Raw contact frequencies for union loops were obtained by intersecting the filtered read pairs from the 11 iPSC and 13 iPSC-CM Hi-C samples with the union loop set using pgltools. These raw contact frequencies were used as input in edgeR^44^, normalized using trimmed mean of M-values (TMM), and compared between the 11 iPSC and 13 iPSC-CM samples using quasilikelihood F-test. The significant differential loops were determined by FDR adjusted q < 0.01.

### Creation of null loop sets for functional comparisons

As chromatin loops, and genome annotations such as chromatin states, are highly structured and depend on genomic distance both between their own anchors and other chromatin loops, we used permutation to test for functional enrichment within chromatin loops and at loop anchors. We generated 1000 null loop sets for both the iPSC called and iPSC-CM called loop sets to use for statistical analysis, as genome-wide background levels of genomic traits may not accurately represent a true random distribution of paired-genomic loci. The null loops were generated for each chromosome by: 1) removing the gap regions on the human reference genome obtained from UCSC genome browser (https://genome.ucsc.edu/) and updating the loop positions according to this “no-gap-genome”; 2) sliding the loop positions on the “no-gap-genome” for a consistent random distance d such that 2Mb < d < chromosome size - 2Mb for each null set; and 3) gap regions were added back to the genome, null loop positions were updated back to hg19. In step 2, when loop positions moved beyond the chromosome size after rotation, loops were instead moved to the beginning of the chromosome. Null loops with anchors overlapping a gap region were removed (an average of 0.5% loops were removed in each cell type).

### Distribution of normalized H3K27ac ChIP-seq and ATAC-seq tag frequencies, and CTCF motifs at anchors

The findMotifsGenome.pl script from HOMER (v4.7) was used to determine enriched motifs at loop anchors, using the entire size of the anchor as the search space. To identify the distribution frequencies of a given motif, or of H3K27ac ChIP-seq and ATAC-seq reads, annotatePeaks.pl with bin size of 500bp and window size of 50kb was used for each set of loops.

### Determining enrichment of chromatin states at loop anchors

For each of the ROADMAP tissues^45^, the core 15-chromatin-state models were obtained as BED format from http://egg2.wustl.edu/roadmap/web_portal/chr_state_learning.html#core_15state, and the states were separated into their original 200bp bins. To determine the enrichment of each chromatin state at a loop anchor, we compared the proportion of 200bp bins in the state of interest on the loop anchor, to the genome-wide background level of the bins via Fisher’s exact test. A significance level of p < (0.05 / 15) was considered significant.

### Identification of differential H3K27ac peaks and differentially expressed genes

To identify differential H3K27ac peaks and genes, we first used featureCounts^46^ to get the number of reads for each assay from each gene from gencode v19, or from each peak identified by merging all the H3K27ac data together. Next, we used DEseq2 v1.10.1^47^ with default parameters to identify differential peaks and genes with a log2(fold change) >2 and an FDR corrected q-value < 0.05.

### Enrichment of cell type specific regulatory regions at CTALs

To determine if cell type specific regulatory regions were enriched at CTALs, for each cell type, we first split the union loop set into CTALs and non-CTALs. Next, we examined whether the proportion of CTALs overlapping a cell type specific regulatory region was statistically larger than the proportion of non-CTALs. For example, to test whether iPSC-CTALs were more likely to harbor an iPSC-specific active promoter, we restricted the analysis to loops overlapping an iPSC active promoter, and tested whether the proportion of loops overlapping an iPSC specific active promoter was higher within CTALs than non-CTALs. For all analyses, we used Roadmap E020 (iPSC) for iPSCs, and Roadmap E083 (fetal heart) for iPSC-CMs. We define an anchor as overlapping a cell type specific regulatory region as an anchor which overlaps the region in the tested cell type (E020 for iPSC-CTALs and E083 for CM-CTALs), but does not overlap the region in the other cell type (E083 for iPSC-CTAL comparisons, E020 for CM-CTAL comparisons).

### Phasing genomes

To obtain accurately phased genotypes for each sample, we performed initial phasing using the Hi-C data, and then subsequently utilized family structure to identify, and fix or remove, haplotyping errors (**point errors**). We first determined the initial phased genotypes for each individual, at each site at least one individual was heterozygous, by analyzing the HiC data with Haploseq^48^. Next, as Haploseq only identifies heterozygous sites, we filled in missing genotype data with unphased genotypes from iPSCORE WGS variant calls for these individuals (Figure S2A). To determine the corresponding parental haplotype for each child haplotype (**parent-child haplotype combination**), we identified the average concordance between each child haplotype, and each of the four parental haplotypes, in 1MB bins chromosome by chromosome, and identified the best matching parent-child haplotype combination for each child chromosome. Within each parent-child haplotype combination, we identified meiotic recombinations within the parent so that we could identify and fix point errors across the genome (Figure S2B). We identified recombinations by finding the extreme points from the following scoring function: for a given child haplotype C1, haplotypes from a single parent PH1 and PH2, and N heterozygotic sites across the genome in the child,

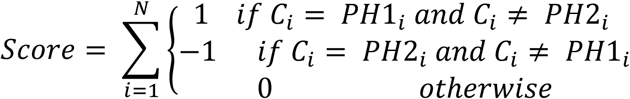

We then split each parent-child haplotype combination into crossover blocks at each crossover position so that each child SNV could be compared to both matching parental haplotypes simultaneously, and fixed switch errors according to Mendelian inheritance. Additionally, if any member of the family was unphased at the site, we phased these variants to follow Mendelian inheritance, generating switch error free genotypes (Figure S2C). After phasing each trio individually, we re-evaluated Mendelian inheritance across all seven individuals, and removed any sites where Mendelian inheritance was violated, as these indicated genotyping errors in one or more individuals.

### Identification of genome-wide imbalanced chromatin loops

To identify Imbalanced Chromatin Loops (**ICLs**), we phased contacts from each chromatin loop in the union loop set across cell types, and identified allelic imbalance that was statistically significant at a genome wide threshold. We first identified all contacts within 25kb of a loop, kept those containing at least one heterozygous SNV, and discarded those with no heterozygous SNVs. We then assigned contacts to their matching haplotype when all heterozygous SNVs matched a single haplotype, and discarded other contacts. Next, at each loop, we calculate a Z score via a binomial approximation to a normal distribution from the major and minor allele counts, and then calculated a p-value from a half-normal distribution for each person. To obtain a single p-value for imbalance of each loop, we use Fisher’s method to obtain a meta-p-value across all 7 individuals. Finally, to identify genome-wide significant ICLs, we use the Benjamini-Hochberg FDR correction to obtain a q-value, and identified loops with a q-value < 0.05 as genome-wide significant ICLs.

### Calculation of power to detect ICLs

To determine the power to identify chromatin loop imbalance at different allelic imbalance fractions, we calculated Z scores as above using parameters for numbers of contacts (ranging from 5-100 in steps of 5), allelic imbalance fractions (from 0.55-0.95 in steps of 0.05). We then calculated the power from a half-normal distribution using alpha thresholds ranging from 1×10^−x^ to 9×10^−x^ for any integer 2 ≤ x ≤ 6 within each individual. We then calculated the alpha threshold from a meta p-value obtained from combining seven individuals displaying the same imbalance via Fisher’s method.

### ASE gene and peak identification

To identify genes and peaks exhibiting genome wide significant allele-specific expression (ASE) from RNA-seq or ChIP-seq data, within each cell type, for each individual, we pooled all samples by cell type, applied WASP^49^ to reduce reference allele mapping bias, and then used MBASED^24^ (R package version 1.4.0) to obtain allelic ratios and p-values for each gene and peak for each individual, and identified significant genes or peaks as those with an FDR corrected q-value < 0.05.

### Chromatin loop set enrichment, and genomic feature enrichment, for ICLs

To identify chromatin loops containing imprinted genes or CNVs, we utilized the pgltools “findLoops” function to create a bed file from the union loop set, and then used bedtools^50^ “intersect” function to obtain all loops containing the genomic characteristic. To identify ASE genes overlapping chromatin loop anchors, we utilized pgltools “intersectlD” function. To identify eQTLs polymorphic in the family with eGenes connected by a chromatin loop, we created a set of all eQTL-eGene pairs with empirical p < 0.05 from DeBoever *et al.*^*21*^ in the PGL format, and utilized pgltools “intersect” to find loops within 20kb of the eQTL-eGene pair. For each genomic feature, we performed a Fisher’s exact test across multiple chromatin loop imbalance p-value thresholds to determine if the genomic feature was enriched in ICLs over the union loop set. To obtain a p-value threshold ICL set, we filtered all chromatin loops to those exhibiting allelic imbalance with a p-value less than or equal to the threshold.

### Determining concordance between loop and molecular phenotype imbalance

To examine the relationship between molecular phenotype (RNA-seq and H3K27ac ChIP-seq) allelic imbalance and chromatin loop imbalance, we compared allelic differences in molecular phenotype data to chromatin loop imbalance frequencies in iPSC-CM data. We first removed chromatin loops containing imprinted genes or CNVs. Next, for each union chromatin loop, we utilized the aforementioned allelic imbalance data; for each molecular phenotype, we pooled the iPSC-CM reads from all samples for each individual, applied WASP^49^ to reduce reference allele mapping bias, and used MBASED to obtain major allele frequencies of each gene/peak. We then identified the most imbalanced SNV in each gene/peak, and used the SNV’s phase to determine the maternal allele frequency of the gene/peak. We then converted maternal allele frequencies to fold changes by dividing the maternal allele frequency by the paternal allele frequency for both molecular phenotypes, and the chromatin loop data.

